# Unraveling the spatial distribution of CAF subsets in PDAC spheroids through a novel spatial flow cytometry approach

**DOI:** 10.1101/2025.08.08.668982

**Authors:** Sara E. Castro-Silva, Camila C. O. M. Bonaldo, Patricia V. B. Palma, Pedro Luiz Porfirio Xavier, Raphael Lucas More, Maristela D Orellana, Sâmia R. Caruso, Rodrigo A. Panepucci

**Affiliations:** Department of Genetics – School of Medicine of Ribeirão Preto (FMRP-USP). Functional Biology Laboratory – Blood Center of Ribeirão Preto; Blood Center – Ribeirão Preto; Laboratory of Comparative and Translational Oncology – Faculty of Animal Science and Food Engineering, University of São Paulo (FZEA-USP); Functional Biology Laboratory – Blood Center of Ribeirão Preto

## Abstract

Pancreatic ductal adenocarcinoma (PDAC) is characterized by a dense stromal compartment, predominantly composed of cancer-associated fibroblasts (CAFs), that contributes to immune exclusion and therapeutic resistance. Although the phenotypic and functional diversity of stromal cells within the TME is well characterized, their spatial distribution and the mechanisms driving this heterogeneity have not been thoroughly investigated. Here, we present SpheroMap Cytometry, an innovative spatial flow cytometry method that enables high-resolution analysis of spatially organized cellular phenotypes within spheroids. CAPAN-1 pancreatic tumor cells and either HS-5 bone marrow stromal or umbilical cord-derived mesenchymal stromal cells (UC-MSCs) were co-cultured in ultra-low-adhesion 96-well plates. After 48 hours aggregation, spheroids were incubated with Image-iT Green Hypoxia to mark hypoxic cells, and after 72 hours spheroids were dissociated and stained with antibodies against CD73 and CD140B. SpheroMap Cytometry revealed that the hypoxic core was significantly enriched for CD73^+^ and myCAF-like populations (CD140B^+^CD73^high^), indicating a functional link between hypoxia and this CAF subpopulation. Moreover, while hypoxia alone was sufficient to drive myCAF differentiation in heterotypic and monotypic spheroids, we found that normoxic induction of myCAF occurred only in the presence of tumor cells supporting the hypothesis that proximity to tumor cells synergizes with hypoxia to regulate CAF differentiation. Our findings demonstrate that hypoxia drives a distinct stromal architecture in PDAC. SpheroMap Cytometry provides a scalable, high-resolution method to dissect the spatial immunophenotype of 3D tumor models, overcoming limitations of static imaging and conventional flow cytometry, opening new avenues for preclinical assessment of stroma-targeting therapies and the development of immunotherapeutics that reprogram the TME.

## 1 Introduction

Pancreatic ductal adenocarcinoma (PDAC) carries one of the worst prognoses among solid malignancies due to nonspecific symptoms, late diagnoses, and therapy resistance, contributing to a five-year survival rate below 11% ^1^. The characteristically immune-excluded profile of PDAC and its consequent resistance to immune checkpoint inhibitor therapies (ICIs) and adoptive cell therapies (ACTs) are attributed to its dense stromal compartment, notably composed of tumor/cancer-associated fibroblasts (TAFs/CAFs) ^2–4^. Within the tumor microenvironment (TME), CAFs constitute the main stromal component, accounting for approximately 80% of the total tumor mass. These populations play a critical role in tumor progression and therapy resistance through their involvement in desmoplasia, immunosuppression, and secretion of factors that promote tumor cell proliferation and survival ^3^.

CAFs coexist as two mutually exclusive and reversible subtypes: FAP^+^αSMA^high myofibroblastic CAFs (myCAFs), which express TGF-β response genes and reside in close proximity to neoplastic cells, receiving juxtacrine tumor-cell signaling; and inflammatory CAFs (iCAFs; αSMA^low^IL-6^high^), which are driven by paracrine signals from tumor cells and secrete inflammatory cytokines such as IL-6, IL-11 and LIF, as well as chemokines including CXCL1 and CXCL2. ^5^. CAF populations with transcriptional, phenotypic and functional profiles analogous to iCAFs and myCAFs are recurrently observed across various cancer types, including PDAC ^5–7^. These two subpopulations express distinct, minimally overlapping marker panels: iCAFs are generally characterized by high levels of CD140A/PDGFRα^High^, CD146^High^, IL6^High^, CXCL12^High^, CD34^High^; whereas myCAFs are defined by high expression of αSMA^High^, FAP^High^, FSP1^High^, CD26^High^, CD140B/PDGFRβ^Med-High^) ^5–9^.

Hypoxia plays a well-established role in the reprogramming and activation of CAFs by promoting tumor-cell secretion of TGF-β and PDGF, thereby favoring the differentiation of precursor cells into myCAFs ^5, 10–14^. It also triggers critical pathways via activation of the transcription factor HIF-1α, which induces transcription of the ecto-enzymes CD39 and CD73 (a key marker of mesenchymal stem cells, MSCs), converting extracellular ATP into adenosine^15–17^.

Although the phenotypic and functional diversity of stromal cells within the TME is well characterized ^2, 18–23^, their spatial distribution and the mechanisms driving this heterogeneity have not been thoroughly investigated. Importantly, the contribution of hypoxia and adenosinergic signaling in this context has not yet been fully elucidated due to the lack of models capable of recapitulating cellular organization within tumor compartments.

In light of these challenges, this study proposes to evaluate how hypoxia influences the distribution of stromal (CAF) cells in spheroid models, through a novel Spatial Flow Cytometry approach (SpheroMap Cytometry) thereby providing a significant methodological advance for the field of translational immuno-oncology.

## 2 Methodology

### 2.1 Cell Lines

CAPAN-1 pancreatic cancer cells (BCRJ, 265) were kindly provided by Dr. Pedro Luiz Porfirio Xavier, collaborating researcher in the Laboratory of Comparative and Translational Oncology (LOCT) at FZEA/USP. HS-5 immortalized stromal cells (ATCC CRL-3611) and umbilical-cord-derived mesenchymal cells (UC-MSCS) were generously supplied by Dr. Sara Teresinha Olalla Saad, Full Professor at the State University of Campinas, and Dr. Maristela Delgado Orellana, researcher at the Hemocenter of Ribeirão Preto, respectively.

### 2.2 Culture and spheroid establishment

The CAPAN-1, HS-5, and primary UC-MSCS cell lines were cultured in IMDM, DMEM, and α-MEM, respectively, each supplemented with 10% fetal bovine serum, 10% penicillin/streptomycin, and HEPES. Cultures were maintained in a 37 °C incubator with 5% CO^+^ and 85% humidity. For spheroid formation total of 20,000 CAPAN-1 and HS-5 or CAPAN-1 and UC-MSCS cells at a 1:4 tumor-to-stromal/mesenchymal ratio were co-plated in ultra-low-adhesion (ULA) 96-well plates (Thermo Scientific 174115), centrifuged and incubated.

### 2.3 Spatial Flow Cytometry (SpheroMap Cytometry)

Given the limitations of conventional flow cytometry and microscopy approaches, we developed a novel spatial flow cytometry method (SpheroMap Cytometry) to enable more detailed immunophenotypic and spatial analysis of the different cell populations within spheroids. Our approach is based on the unique properties of the Image-iT Green Hypoxia reagent, which accumulates intracellular fluorescence specifically in cells located within the hypoxic core of the spheroid. Importantly, this fluorescent signal is retained even after enzymatic dissociation of the spheroids, thus allowing for quantitative immunophenotypic analysis of the spatial distribution of distinct cellular subpopulations within the hypoxic (central) and normoxic (peripheral) regions of the spheroid.

#### 2.3.1 Distribution of CAF subpopulations

In summary, only CAPAN-1 tumor cells were pre-labeled with Cell Proliferation Dye eFluor 450 (at a final concentration of 10 µM), then co-seeded with stromal/mesenchymal cells and incubated. After 48 hours, half of the medium was replaced with medium containing Image-iT Green Hypoxia (Invitrogen™ I14833, at a final concentration of 1.25 µM), and on the following day, spheroids were collected, transferred to 15 mL centrifuge tubes (8 spheroids per tube, totaling 160,000 cells), washed, and incubated in 2 mL of 0.05% Trypsin-EDTA (Gibco 15400054) for 10 to 15 minutes for enzymatic and mechanical dissociation (using a pipette with a 200 µL tip). The dissociated cells were then washed with RPMI medium, centrifuged, and resuspended in PBS + 2% FBS, followed by incubation for 30 minutes at 4 °C with PE Mouse Anti-Human CD140B antibody (Clone 28D4, BD Pharmingen™ 558821) to identify the myCAF population within distinct spheroid compartments, and APC Mouse Anti-Human CD73 antibody (Clone AD2, BD Pharmingen™ 560847) to detect CD73 (NT5E) expression in these compartments.

### 2.4 Analysis of SpheroMap Cytometry Data

SpheroMap Cytometry data were acquired using a BD FACSLyric™ Clinical Flow Cytometry System, then exported and analyzed using FlowJo software v.10.9.0. Data obtained in FlowJo were subsequently exported to GraphPad Prism 9 for graph generation and statistical analysis. The statistical test used was ordinary two-way ANOVA followed by Sidak’s multiple comparisons test.

## 3 Results

### 3.1 Distribution of CAF Subpopulations in spheroid compartments

Automated quantitative fluorescence microscopy (High Content Screening-HCS, ImageXpress Micro XLS; Molecular Devices) (Fig. 2A) revealed that the experimental condition (20,000 cells at a 1:4 ratio) enabled the formation of well-compacted spheroids as early as 24 hours in co-cultures of CAPAN-1 and HS-5 cells. In contrast, spheroids formed with MSCs exhibited loose aggregation, with MSCs not yet fully incorporated into the spheroid structure at this time point. However, from 48 hours onward, we observed more compact spheroids.

**Fig. 1.**
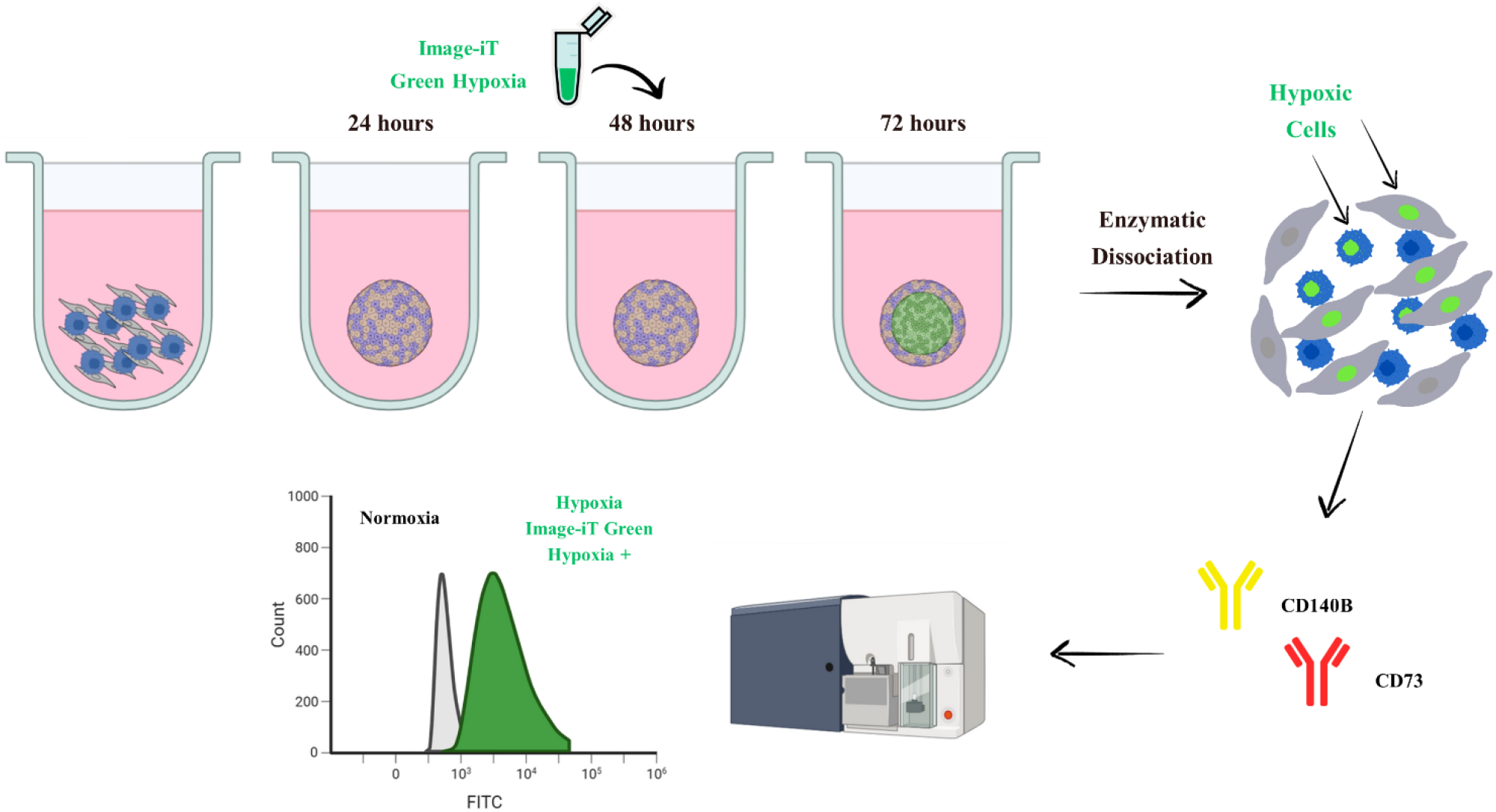
Schematic of the SpheroMap Cytometry methodology. A total of 20,000 tumor cells (pre-labeled) were co-plated in U-bottom ultra-low attachment plates with unlabeled stromal/mesenchymal cells at a 1:4 tumor-to-stromal ratio. 48 hours after seeding, the Image-iT Green Hypoxia (1.25 µM) reagent was added to wells containing the spheroids. Intracellular fluorescence accumulates in cells occupying the hypoxic core of the spheroid and, importantly, is retained even after enzymatic dissociation, enabling immunophenotypic analysis of cells that inhabited the hypoxic region. 72 after initial seeding, spheroids were enzymatically dissociated, stained with antibodies against myCAF (CD140B) and CD73, and subjected to analysis.

**Fig. 2.**
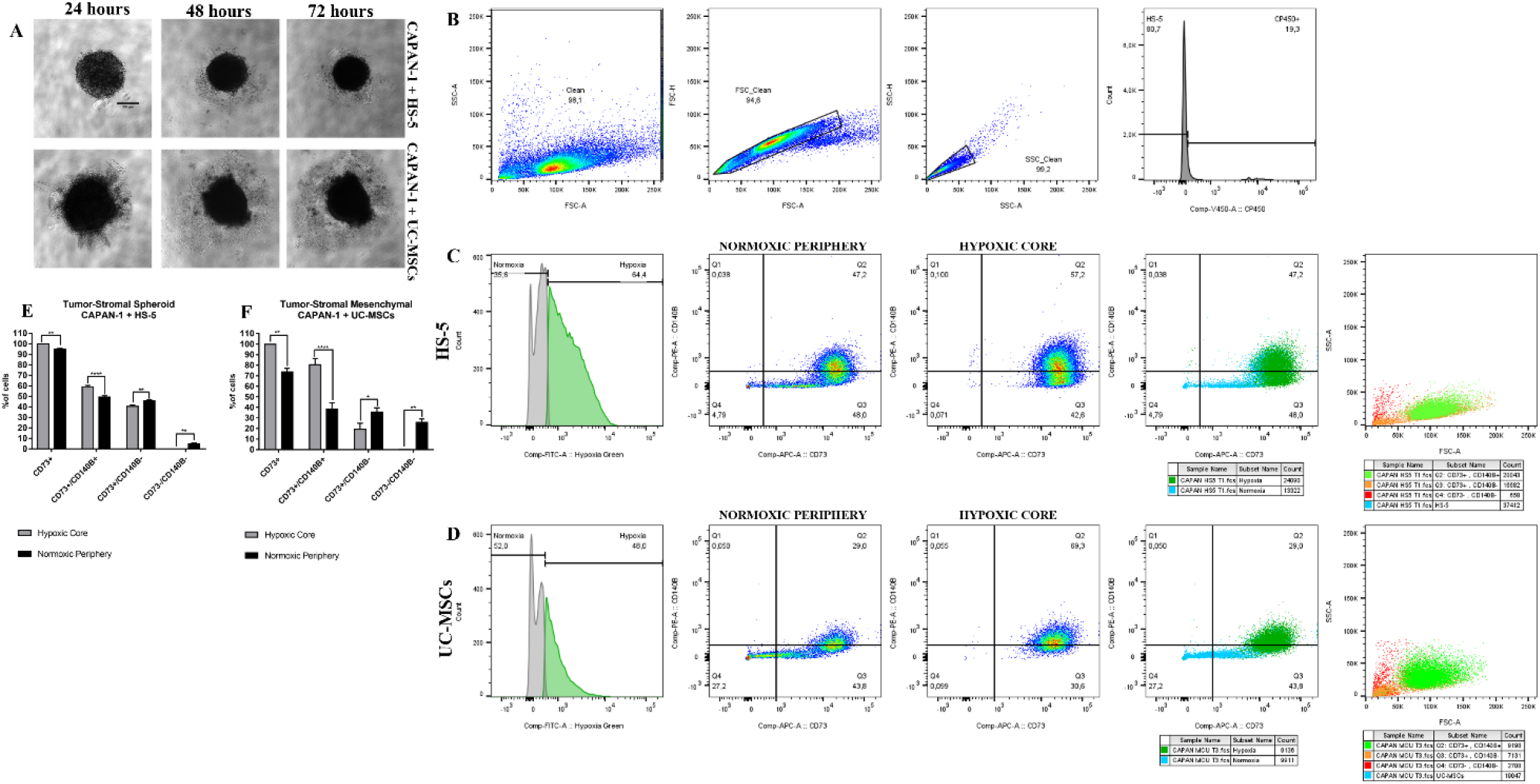
Expression of CD73 and CD140B in stromal/mesenchymal cells from heterotypic spheroids. Spheroids were formed by co-culturing tumor cells (pre-labeled with Cell Proliferation Dye eFluor 450; CP450, V450 channel) and stromal/mesenchymal cells at a 1:4 ratio, with a total of 20,000 cells, for 72 hours post-seeding. The day before analysis (72 hours post-seeding), the hypoxia reagent (Image-iT Green Hypoxia; FITC) was added to the spheroid wells at a final concentration of 1.25 µM to label cells residing in the hypoxic environment. The following day, spheroids were enzymatically dissociated, incubated with anti-CD140B (PE) and anti-CD73 (APC) antibodies, and subjected to flow cytometric analysis. **(A)** Spheroids were imaged after 24, 48, and 72 hours using the ImageXpress XLS HCS system with a 4X objective, transmitted light (phase contrast, PhL). **(B)** The gating strategy involved sequential steps to ensure accurate identification of distinct cellular subpopulations. Debris and dead cells were first excluded based on FSC-A vs. SSC-A gating, followed by singlet discrimination using FSC-H vs. FSC-A and SSC-H vs. SSC-A plots. Next, using CP450 and DAPI staining, tumor cells and dead events (CP450^+^) were separated from stromal/mesenchymal cells (CP450^-^). **(C-D)** The gates shown were applied in the following sequence: hypoxic and non-hypoxic cells were identified based on Image-iT Green staining. A subsequent gating step classified these cells as CD140B^+^, CD73^+^, double-positive or double-negative. Next, using the FSC/SSC parameters on the total HS-5/UC-MSCS cell population, we plotted all CD73^-^/CD140B^-^, CD73^+^/CD140B^-^, and CD73^+^/CD140B^+^ cells to correlate cell size with marker expression. **(E-F)** Statistical analysis was performed using ordinary two-way ANOVA followed by Sidak’s multiple comparisons test. *p < 0.05, **p < 0.01, ****p < 0.0001.

For a given cell type, the intensity of Image-It Green is proportional to the level of hypoxia, resulting in a radial distribution, with cells in the inner core showing higher fluorescence intensities. Thus, the intensity of Image-It Green defines a functional compartment of hypoxic cells confined to the inner central region of the spheroid. However, since different cell types can display different staining intensities, it is important to notice that the spatial core region functionally-defined by Image-It Green staining (i.e. the hypoxic compartment) is specific for a given cell type, as for a different cell type, the physical space occupied by the hypoxic cell may differ. Thus, throughout this manuscript we use the term “hypoxic compartment” to refer to a spatially-defined core region, only in the context of comparison made for a given cell type.

In spheroids formed with HS-5 cells (Fig. 2C), CD73 positivity was high in both compartments, though slightly higher in the hypoxic core compared to the normoxic periphery (99.83% vs. 95.00%). CD140B expression also varied between compartments, being higher in the hypoxic core than in the periphery (81.00% vs. 68.63%), with a greater proportion of CD140B^-^ cells observed in the periphery (31.37% vs. 19.00%). These differences were reflected in the co-expression of CD73^+^/CD140B^+^, which was higher in cells located in the hypoxic core compared to the normoxic periphery (59.27% vs. 49.37%) (Fig. 2E).

The results observed in spheroids formed with UC-MSCS cells were similar (Fig. 2D). The hypoxic core showed a higher proportion of CD73^+^ cells compared to the normoxic periphery (99.83% vs. 73.67%). CD140B expression also differed between compartments, being higher in the hypoxic core than in the periphery (82.53% vs. 55.87%), with a greater proportion of CD140B^-^ cells in the periphery (44.13% vs. 17.47%). These findings were reflected in the co-expression of CD73^+^/CD140B^+^, which was greater in cells located in the hypoxic core compared to the normoxic periphery (80.57% vs. 38.50%) (Fig. 2F). When we analyzed the FSC/SSC parameters on the total HS-5/UC-MSCS cell population to correlate cell size with marker expression, we observed that as cells increased in size, they began to express both markers (CD73^+^/CD140B^+^) (Fig. 2C-D).

### 3.2 Expression of CD140B/CD73 in monotypic spheroids

We observed that the stromal/mesenchymal cells appeared to comprise two populations with distinct morphologies, one “small” and one “large.” To rule out the possibility that the smaller population was actually tumor cells that had lost the Cell Proliferation Dye eFluor 450 labeling, spheroids composed solely of stromal or mesenchymal cells—while maintaining the total count of 20,000 cells—were formed to confirm the morphological profile of the stromal/mesenchymal cells.

The expression of the myCAF phenotype in the hypoxic region of spheroids composed solely of stromal (~40%) (Fig. 3B) or mesenchymal (~45%) (Fig. 3C) cells was similar to that observed in the hypoxic region of tumor–stromal/mesenchymal co-culture spheroids (Fig. 2C-D), indicating that the myCAF immunophenotype is strongly regulated by hypoxia. However, the normoxic regions of both mono- and heterotypic spheroids diverged: in heterotypic spheroids myCAF expression was approximately 47% in tumor–stromal (Fig. 2B) and 30% in tumor– mesenchymal (Fig. 2C) spheroids. In contrast, in monotopic spheroids myCAF expression dropped dramatically in the normoxic region, to about 10% in stromal-only (Fig. 3B) and roughly 1% in mesenchymal-only spheroids (Fig. 3C).

**Fig. 3.**
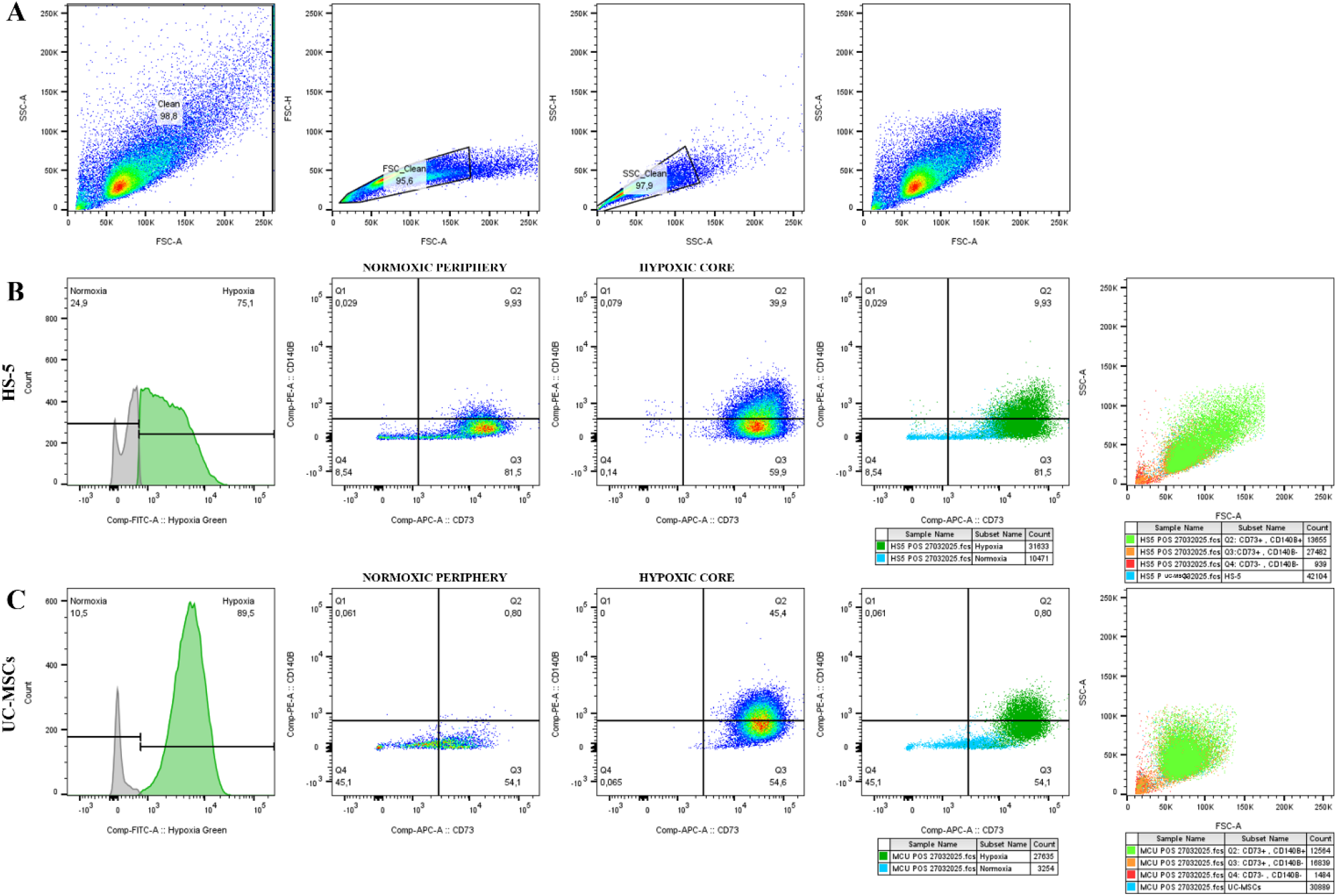
Expression of CD73 and CD140B in stromal/mesenchymal cells from monotypic spheroids. Stromal and mesenchymal spheroids were formed with a total of 20,000 cells, for 72 hours post-seeding. On the day before analysis (72 hours post-seeding), the hypoxia reagent (Image-iT Green Hypoxia; FITC) was added to the spheroid wells at a final concentration of 1.25 µM to label cells residing in the hypoxic environment. The following day, spheroids were enzymatically dissociated, incubated with anti-CD140B (PE) and anti-CD73 (APC) antibodies, and subjected to flow cytometric analysis. **(A)** The gating strategy involved sequential steps to ensure accurate identification of distinct cellular subpopulations. Debris and dead cells were first excluded based on FSC-A vs. SSC-A gating, followed by singlet discrimination using FSC-H vs. FSC-A and SSC-H vs. SSC-A plots. **(B-C)** The gates shown were applied in the following sequence: hypoxic and non-hypoxic cells were identified based on Image-iT Green staining. A subsequent gating step classified these cells as CD140B^+^, CD73^+^, double-positive or double-negative. Next, using the FSC/SSC parameters on the total HS-5/UC-MSCS cell population, we plotted all CD73^-^/CD140B^-^, CD73^+^/CD140B^-^, and CD73^+^/CD140B^+^ cells to correlate cell size with marker expression.

The results obtained reveal that, in hypoxic environments, myCAF profile expression occurs regardless of co-culture with tumor cells, indicating a critical role for hypoxia; however, in the absence of hypoxia, the myCAF profile is present only in heterotypic spheroids, indicating that its expression is regulated by the proximity of stromal cells to tumor cells. Again, when we analyzed the FSC/SSC parameters on the total HS-5/UC-MSCS cell population, we observed that as cells increased in size they began to express both markers (CD73^+^/CD140B^+^).

### 3.3 Expression of CD140B/CD73 in stromal/mesenchymal populations FSC^low^/SSC^low^and FSC^high^/SSC^high^ from heterotypic spheroids

Once we confirmed that the cells we were analyzing were indeed only stromal/mesenchymal cells, we decided to separate these two populations based on their morphology and analyze CD73 and CD140B expressions in each. We designated the “large” population as FSC^high^/SSC^high^ and the “small” population as FSC^low^/SSC^low^.

The results obtained for both CAPAN-1/HS-5 and CAPAN-1/UC-MSCS heterotypic spheroids were similar. In the hypoxic core, the FSC^high^/SSC^high^ subpopulation predominated over the FSC^low^/SSC^low^ subpopulation (70% vs. 3%). Both subpopulations were positive for CD73 at nearly 100%, but they differed in CD140B expression, which was higher in the FSC^high^/SSC^high^ subpopulation compared to the FSC^low^/SSC^low^ subpopulation (80% vs. 9%). The FSC^low^/SSC^low^ group showed a higher percentage of CD140B-negative cells (90% vs. 22%), which was reflected in the co-expression of CD73^+^/CD140B^+^: 60% for FSC^high^/SSC^high^ versus 5% for FSC^low^/SSC^low^ (Fig 4B-E). When we analyzed the normoxic environment, we observed that the subpopulation results were inverted: the FSC^low^/SSC^low^ subpopulation predominated over the FSC^high^/SSC^high^ subpopulation (98% vs. 33%). Both subpopulations showed high CD73 positivity, with FSC^high^/SSC^high^ still predominating (99% vs. 82%). CD140B expression also varied: FSC^high^/SSC^high^ had a greater proportion of CD140B-positive cells than FSC^low^/SSC^low^ (85% vs. 2%), while FSC^low^/SSC^low^ had more CD140B-negative cells (99% vs. 15%). These results were mirrored in the co-expression of CD73^+^/CD140B^+^, which was higher in the FSC^high^/SSC^high^ subpopulation than in the FSC^low^/SSC^low^ subpopulation (63% vs. 1%) (Fig 4B-E).

**Fig. 4.**
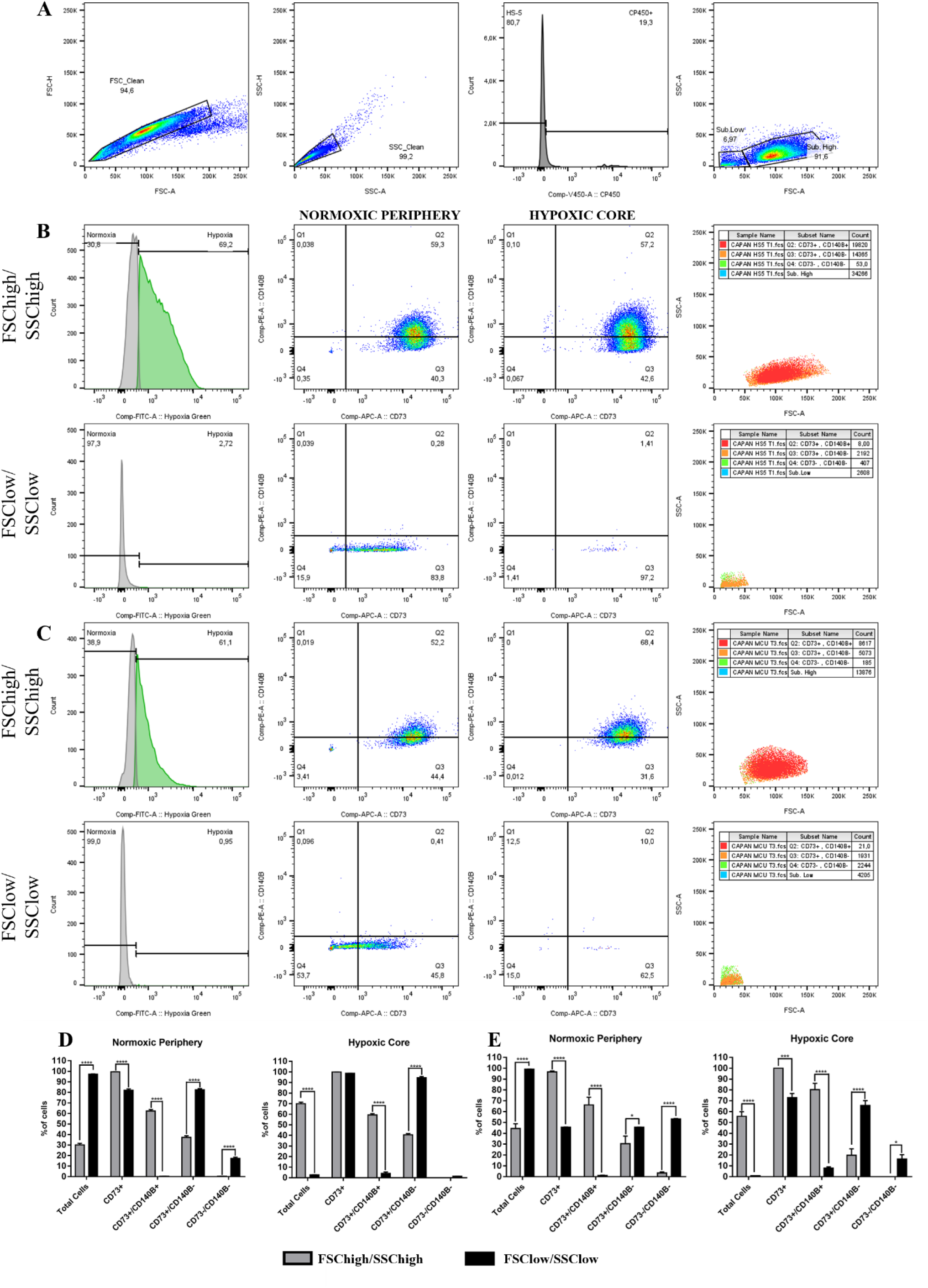
Expression of CD73 and CD140B in FSC^high^/SSC^high^ and FSC^low^/SSC^low^ stromal/mesenchymal cells from heterotypic spheroids. Spheroids were formed by co-culturing tumor cells (pre-labeled with Cell Proliferation Dye eFluor 450; CP450, V450 channel) and stromal/mesenchymal cells at a 1:4 ratio, with a total of 20,000 cells, for 72 hours post-seeding. The day before analysis (72 hours post-seeding), the hypoxia reagent (Image-iT Green Hypoxia; FITC) was added to the spheroid wells at a final concentration of 1.25 µM to label cells residing in the hypoxic environment. The following day, spheroids were enzymatically dissociated, incubated with anti-CD140B (PE) and anti-CD73 (APC) antibodies, and subjected to flow cytometric analysis. **(A)** The gating strategy involved sequential steps to ensure accurate identification of distinct cellular subpopulations. Debris and dead cells were first excluded based on FSC-A vs. SSC-A gating, followed by singlet discrimination using FSC-H vs. FSC-A and SSC-H vs. SSC-A plots. Next, using CP450 and DAPI staining, tumor cells and dead events (CP450^+^) were separated from stromal/mesenchymal cells (CP450^-^) and a second gating step was performed to separate the FSC^high^/SSC^high^ and FSC^low^/SSC^low^ populations. **(B-C)** In the FSC^high^/SSC^high^ and FSC^low^/SSC^low^ subpopultion of HS-5/UC-MSCS cells, the gates shown were applied in the following sequence: hypoxic and non-hypoxic cells were identified based on Image-iT Green staining. A subsequent gating step classified these cells as CD140B^+^, CD73^+^, double-positive or double-negative. Next, using the FSC/SSC parameters on the total HS-5/UC-MSCS cell population, we plotted all CD73^-^/CD140B^-^, CD73^+^/CD140B^-^, and CD73^+^/CD140B^+^ cells to correlate cell size with marker expression **(D-E)** Statistical analysis was performed using ordinary two-way ANOVA followed by Sidak’s multiple comparisons test. *p < 0.05, ***p < 0.001, ****p < 0.0001.

## 4 Discussion

The stromal component of PDA comprises approximately 80–90% of the total tumor mass, playing a key role in establishing physical and functional barriers that limit immune cell access to neoplastic cells, and thus being crucial for therapy resistance. Notably, CAF populations contribute significantly to tumor progression and therapy resistance through their involvement in desmoplasia, immunosuppression, and secretion of factors that promote cancer cell proliferation and survival ^3, 24^. As a consequence of tumorigenesis, hypoxia and inflammation within the tumor trigger critical processes through activation of the Hypoxia-Inducible Factor (HIF), a master regulator of tumor progression. Hypoxia is known to drive CAF reprogramming and activation by promoting tumor-cell secretion of TGF-β and PDGF, which in turn favor the differentiation of precursor cells into myCAFs. Previous studies have reported that myCAFs constitute approximately 50% of all CAFs, making them the most abundant CAF subpopulation in PDAC stroma; they are also predominantly located adjacent to neoplastic cells, receiving juxtracrine signaling from tumor cells ^5, 10–14^.

Our results revealed that the myCAF marker CD140B was highly expressed in the stromal cells, particularly in the hypoxic region, both in monotypic and heterotypic spheroids clearly linking the hypoxic environment to a myofibroblastic immunophenotypic profile. However, the expression of the myCAF marker in the normoxic environment of only heterotypic spheroids suggests that this profile is also regulated by proximity to tumor cells due to TGF-β secretion by tumor cells. The proximity of myCAFs to tumor cells has already been demonstrated in the literature ^5^.

The concurrent analysis of CD73 and CD140B (myCAF) in stromal cells from both compartments showed that CD73 was also highly expressed in myCAFs in both the hypoxic and normoxic regions, with this dual expression being greater in the hypoxic compartment. HIF-1α activation by hypoxia induces transcription of the ecto-enzymes CD39 and CD73 (a key MSC marker), which convert extracellular ATP into adenosine ^15–17^. CD73 has been reported to be highly expressed in solid tumors, including CAFs in PDAC, where it functions as a major immunosuppressive mediator within the TME ^25–28^.

Our results indicated that the stromal and mesenchymal populations exhibited two size‐based subgroups, which we designated FSC^low^/SSC^low^ and FSChigh/SSChigh. This morphological difference may be associated with CAF activation: when activated—due to hypoxia and cytokine secretion by tumor cells—CAFs adopt a star‐shaped, more elongated morphology, in contrast to quiescent fibroblasts ^13, 29–31^. The fact that FSC^high^/SSC^high^ cells are likely activated explains their greater abundance in hypoxic regions and their higher expression of CD73 and CD140B.

Our findings are consistent with the literature, which not only corroborates its hypoxia-driven regulation but also allowed us to clearly associate the hypoxic environment with the CD140B^+^CD73^high^ stromal immunophenotype linked to putative myCAFs.

## 5 Conclusion

The present study established an innovative, multidimensional experimental platform to systematically investigate how hypoxia and metabolic gradients affect the spatial organization and functional profile of stromal cell populations within the tumor microenvironment of pancreatic ductal adenocarcinoma. The principal methodological innovation of this study, SpheroMap Cytometry, proved to be a powerful and scalable tool for inferring the relative spatial localization of cancer-associated fibroblasts in PDAC spheroids, overcoming the limitations of conventional approaches such as static microscopy and traditional flow cytometry. Using this technique, we were able to identify that the myCAF immunophenotype (CD140B^+^CD73^high^) is associated with the tumor’s hypoxic region; moreover, in addition to hypoxia, the myCAF immunophenotype can also be regulated by the proximity of stromal cells to tumor cells, allowing its expression in normoxic regions and further enhancing its expression in the hypoxic region of heterotypic spheroids.

These findings underscore the central role of tumor architecture and hypoxic gradients in modulating cell behavior within the TME and highlight the importance of approaches that combine physiological relevance with high analytical capability. The platform developed in this study represents a promising tool for preclinical screening of immunotherapies and combination treatments targeting the TME, with potential translational applications in experimental oncology.

## 6 Acknowledgements

This study was funded by the São Paulo Research Foundation (FAPESP) Brasil, process #2022/12856-6; the Brazilian Federal Agency for Support and Evaluation of Graduate Education (CAPES); and the National Council for Scientific and Technological Development (CNPQ).

## Notes

### Competing Interest Statement

The authors have declared no competing interest.

